# No evidence of paralogous loci or new *bona fide* microRNAs in telomere to telomere (T2T) genomic data

**DOI:** 10.1101/2021.12.09.471935

**Authors:** Arun H. Patil, Marc K. Halushka, Bastian Fromm

## Abstract

The telomere to telomere (T2T) genome project discovered and mapped ∼240 million additional base pairs of primarily telomeric and centromeric reads. Much of this sequence was comprised of satellite sequences and large segmental duplications. We evaluated the extent to which human *bona fide* microRNAs (miRNAs) may be found in additional paralogous genomic loci or if previously undescribed microRNAs are present in these newly sequenced regions of the human genome. New genomic regions of the T2T project spanning ∼240 million bp of sequence were obtained and evaluated by blastn for the human miRNAs contained in MirGeneDB2.0 (N=556) and miRBase (N = 1917) along with all species of MirGeneDB2.0 miRNAs (N=10,899). Additionally, bowtie was used to compare unmapped reads from >4,000 primary cell samples to the new T2T sequence. Based on sequence and structure, no *bona fide* miRNAs were identified. Ninety-seven miRNAs of questionable authenticity (frequently known repeat elements) were identified from the miRBase dataset across the newly described regions of the human genome. These 97 represent only 51 miRNA families due to paralogy of highly similar miRNAs such as 24 members of the hsa-mir-548 family. Altogether, this data strongly supports our having identified widely expressed *bona fide* miRNAs in the human genome and move us further toward the completion of human miRNA discovery.

## Introduction

microRNAs (miRNAs) are a class of small regulatory RNAs that block protein translation. Since their discovery in 1993, a large amount of research has sought to determine the number of, and types of members of this RNA family (Lee et al. 1993; Lagos-Quintana et al. 2001; Halushka et al. 2018). This is a contentious issue in science, with a number of different estimates of miRNA family size depending on competing views of the criteria for designating small RNA sequences of on average 22 bp as miRNAs (Griffiths-Jones 2006; Fromm et al. 2015; Backes et al. 2018; Fromm et al. 2020).

While miRBase used to be a repository for published miRNA candidates, where authors could submit their published miRNA candidates (Griffiths-Jones 2004), MirGeneDB uses a uniform system for microRNA annotation & nomenclature, based on next generation sequencing detectable criteria of miRNA biogenesis to arrive at *bona fide* miRNA complements for metazoan species (Fromm et al. 2015; Fromm et al. 2020; Fromm et al. 2021).

Additionally, due to the ancient origins of miRNAs and genomic rearrangements across species, including segmental duplications, several miRNAs are expressed from multiple genomic loci (Peterson et al. 2021). For example, the miRNA hsa-let-7a is found on chromosomes 9, 11, and 22 (Hsa-Let-7-P2a1, Hsa-Let-7-P1a, Hsa-Let-7-P1D), with the -5p (mature) sequence being identical across all three loci. Other miRNAs have undergone minimal sequence changes through these duplication events and now exist as families such as the MIR-17 family, from which hsa-miR-18a (Hsa-Mir-17-P2a) on chromosome 13 and hsa-miR-18b (Hsa-Mir-17-P2c) on the X chromosome differ only by the 20^th^ base of their -5p (dominant) sequence.

A significant amount of our understanding of miRNA expression patterns comes from small RNA-sequencing datasets that capture miRNAs, tRNA halves and fragments, rRNA fragments and other RNA species (de Rie et al. 2017; McCall et al. 2017; Lorenzi et al. 2021; Patil and Halushka 2021). A common curiosity of small RNA-seq data is a significant number of unmapped reads in these datasets. A number of rationales have been ascribed to this material. Some explanations for this extra data include poor sequence quality reads, contamination, infectious organisms, and repeat elements. Another explanation could be that the sequence aligns to the unmapped and uncharacterized regions of the human genome.

Despite the declaration of completing the human genome in 2001, we have always known that large segments of pericentromeric and peritelomeric regions remained unmapped (Lander et al. 2001). Recently, the telomere to telomere (T2T) consortia has used long read sequencing and other methods to fully map a complete human genome (Nurk et al. 2021). This effort increased the amount of sequenced and aligned genome by ∼8%. Most of this sequence is repetitive pericentromeric satellite structures (Altemose et al. 2021; Vollger et al. 2021). However, some segmental duplications and other gene-containing genomic structures exist in this area, which may harbor miRNA paralogs or even heretofore unmapped *bona fide* miRNAs (Nurk et al. 2021; Vollger et al. 2021).

We wondered if this newly characterized region of the human genome contained paralogous loci of known miRNAs and whether any yet described *bona fide* miRNAs may exist in these areas. To do this we surveyed ∼240 million bp of T2T sequence using both known miRNAs and unmapped sequence from 4,150 mostly primary cell type datasets.

## Methods

### Data set data acquisition

The complete T2T genome was obtained from https://github.com/marbl/CHM13 (v1.0) retrieved 7/30/2021) (Nurk et al. 2021). It is also located at https://www.ncbi.nlm.nih.gov/assembly/GCA_009914755.2. Dr. Mitchell Vollger kindly provided a list of non-syntenic regions of T2T relative to GRCh38, which were selected from the full T2T genome using getfasta (BEDTools)(Quinlan 2014). This 240 million bp dataset was subsequently used for all searching (hereafter, the T2T genome). A list of both homo sapiens and “All species” miRNA precursor sequences were obtained from MirGeneDB (https://mirgenedb.org/download, accessed 7/30/21) (Fromm et al. 2020). Separately, the hairpin.fa file from miRBase.org was obtained (https://www.mirbase.org/ftp.shtml accessed 7/30/21) and the homo sapiens (hsa) miRNAs were subsetted out (Griffiths-Jones 2006).

### Unaligned small RNA-seq reads

From an ongoing project to characterize the cellular microRNAome, 4,150 primary cell, plasma, exosome, and cancer cell line small RNA-seq files were obtained from the Sequence Read Archive (SRA) and processed through miRge3.0 using parameters (miRge3.0 -s file_names -gff -bam -trf -lib /miRge3_Lib/ -on human -db mirbase -o <output folder> -mEC -ks 20 -ke 20 -a variable_adapters) (Patil and Halushka 2021). All unmapped reads from the 4,150 files were concatenated and collapsed into a list of unique reads with the read count denoted using a python (3.8.5) script.

### Alignment to the non-syntenic regions of T2T

All reported miRNAs from MirGeneDB and miRBase were used in a local blastn (version: 2.6.0+) search with the non-syntenic T2T sequence. Parameters were: “blastn -query mirgenedb_allSps.fasta -db../blastthis/jhu_chm13_v1.0_T2T.fasta -html > blast_output_preAllSps_miRGeneDB.html”. Unmapped reads from small RNA-seq datasets of length 16-27 bp were aligned to the non-syntenic regions of T2T using Bowtie (v1.2.3) and command line: “bowtie <T2T index> -f all_unmappedReads.fasta -S <Sam Output> -p <threads> -n 0” (Langmead 2010).

## Results

### Search for paralogous miRNA loci in T2T regions

There were 126 separate non-syntenic T2T segments with a total read length of 240,044,315 bp. This was the search space for all further miRNA analyses. These segments represented ∼8% new sequence compared to GRCh38. Both the homo sapiens (N=556) and all species (N=10,899) searches of MirGeneDB (*bona fide*) miRNA precursors yielded zero blastn alignments to the T2T data indicating no previously undiscovered paralogous miRNA loci.

A search of human miRBase miRNA hairpins (N=1917) identified 97 miRNA hairpins with alignments to the T2T data (Table 1). In total, 1794 separate alignments with a percent identity between 80.5 and 100% were detected. There were 445 alignments with a perfect match across a range of 27-179 bases, with roughly half (226), having a perfect alignment of only 27-30 bases. The average length of the hairpin sequence was 82 bp. The dominant miRNA found in this collection was the hsa-miR-548 class. This putative miRNA shares high sequence homology with the Made1 (Tc1/Mariner) repeat family (Piriyapongsa and Jordan 2007). miRbase lists 73 members of the miR-548 family. Twenty-four of these were listed among the miRNAs with alignment in the T2T sequences. Other miRNAs listed repeatedly, such as hsa-miR-1302 (MER53 element), hsa-miR-3118 (Line 1 element), and hsa-miR-5701 (Rep522 repeat), were generally overlapping known repeat elements. The miRNA hairpin with the highest sequence identity was hsa-miR-3648-1, a putative miRNA with a high G/C content (81%), which matched identically (180 bp) in four T2T locations: Two locations in Chr. 13 and one each on Chr. 14 and 21.

**Table 1:**
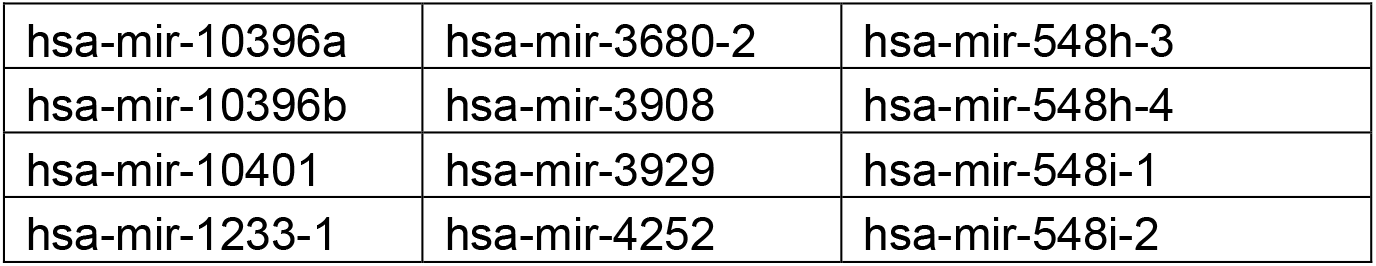
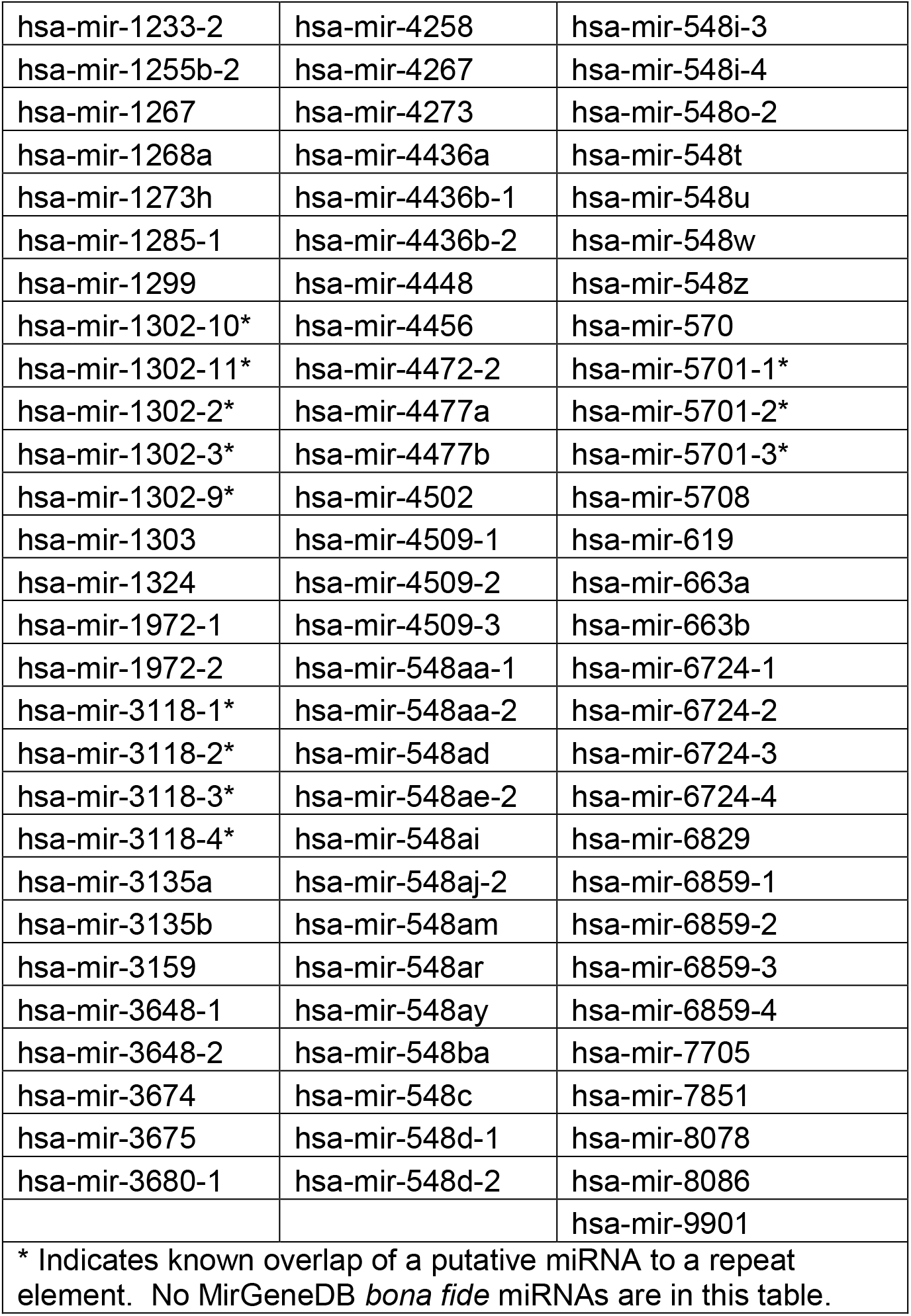
miRBase putative miRNAs aligned to T2T sequence

### Novel miRNA discovery

From 4,150 small RNA-seq datasets of primarily primary cell sequencing that had been processed through miRge3.0, 2,872,614,004 unmapped reads were collapsed into a single search space. Using Bowtie alignment without mismatches, 296,475 unique 16-27bp length reads aligned, representing 383,109,405 total reads mapping to the T2T sequences. These sequences were reviewed for the likelihood they may represent sequences from a miRNA hairpin loop (multiple sequences aligning with the same 5p edge (±1bp) separated from at least one additional sequence by 10-25 bp loop), yielding 8 potential hits. RNA folding of these 8 sequences did not generate a viable stem loop structure. Thus, no novel miRNAs were described.

### Conclusions

A thorough investigation of 240 million bases of new T2T sequences did not identify any new *bona fide* miRNAs or paralogous genomic regions containing known *bona fide* miRNAs. 97 miRNAs of questionable provenance from miRBase were aligning to the T2T sequence, but these most likely represent repeat elements that have been misassigned as miRNAs.

Although every explorer looks for new worlds to conquer, there is some satisfaction in knowing we are nearing the end of the discovery of widely expressed human *bona fide* miRNAs. While it is possible that rare human cell types may harbor miRNAs yet found in RNA-seq, there is no reason to think they will be found in pericentromeric or peritelomeric regions. One last area of potential discovery would be larger genomic regions variable between ethnic populations. It is possible that groups with lower requirements for miRNA discovery may claim findings in the new T2T space, but one should be cautious, if those reports appear.

## Acknowledgements

The authors thank Mitchell Vollger and Evan Eichler for their assistance and helpful comments.

## Funding

A.H.P. and M.K.H. were supported by grants R01HL137811 and R01GM130564. B.F. is supported by the Tromsø forskningsstiftelse (TFS) [20_SG_BF ‘MIRevolution’].

## Notes

### Competing Interest Statement

The authors have declared no competing interest.

